# Scaffold-scaffold interactions regulate cell polarity in a bacterium

**DOI:** 10.1101/2020.08.21.262345

**Authors:** Wei Zhao, Samuel W. Duvall, Kimberly A. Kowallis, Chao Zhang, Dylan T. Tomares, Haley N. Petitjean, W. Seth Childers

**Author notes:** Corresponding Author: W. Seth Childers, Chevron Science Center, Room 801, 219 Parkman Avenue, Pittsburgh, PA 15260. Equal contributions (Wei Zhao, Samuel W. Duvall).

## Abstract

The localization of two biochemically distinct signaling hubs at opposite cell poles provides the foundation for asymmetric cell division in *Caulobacter crescentus*. Here we identify an interaction between the scaffolds PodJ and PopZ that regulates the assembly of the new cell pole signaling complex. Time-course imaging of a mCherry-sfGFP-PopZ fluorescent timer throughout the cell cycle revealed that existing PopZ resides at the old cell pole while newly translated PopZ accumulates at the new cell pole. Our studies suggest that interactions between PodJ and PopZ promotes the sequestration of older PopZ and robust accumulation of newl PopZ at the new cell pole. Elimination of the PodJ-PopZ interaction impacts PopZ client proteins, leading to chromosome segregation defects in one-third of cells. Additionally, this PopZ-PodJ interaction is crucial for anchoring PodJ and preventing PodJ extracellular loss at the old cell pole through unknown mechanism. Therefore, segregation of PopZ protein at the old pole and recruitment of newly translated PopZ at the new pole via the PodJ scaffold ensures stringent inheritance and maintenance of the polarity axis within dividing *C. crescentu*s cells.

## Introduction

Scaffolding proteins can direct and rewire information flow in cellular signaling networks^1^. Through the recruitment of signaling proteins into multi-enzymatic complexes, scaffolding proteins give rise to cellular functions such as cytoskeletal dynamics, cell polarity, division, and morphogenesis^1,2^. In the bacterium *Caulobacter crescentus*, a set of spatiotemporally distributed scaffolding proteins are essential for the establishment and maintenance of cell polarity. This underlying asymmetry enables *Caulobacter crescentus* to divide into a motile swarmer cell and a sessile stalked cell^3-5^ (Figure 1).

**Figure 1:**
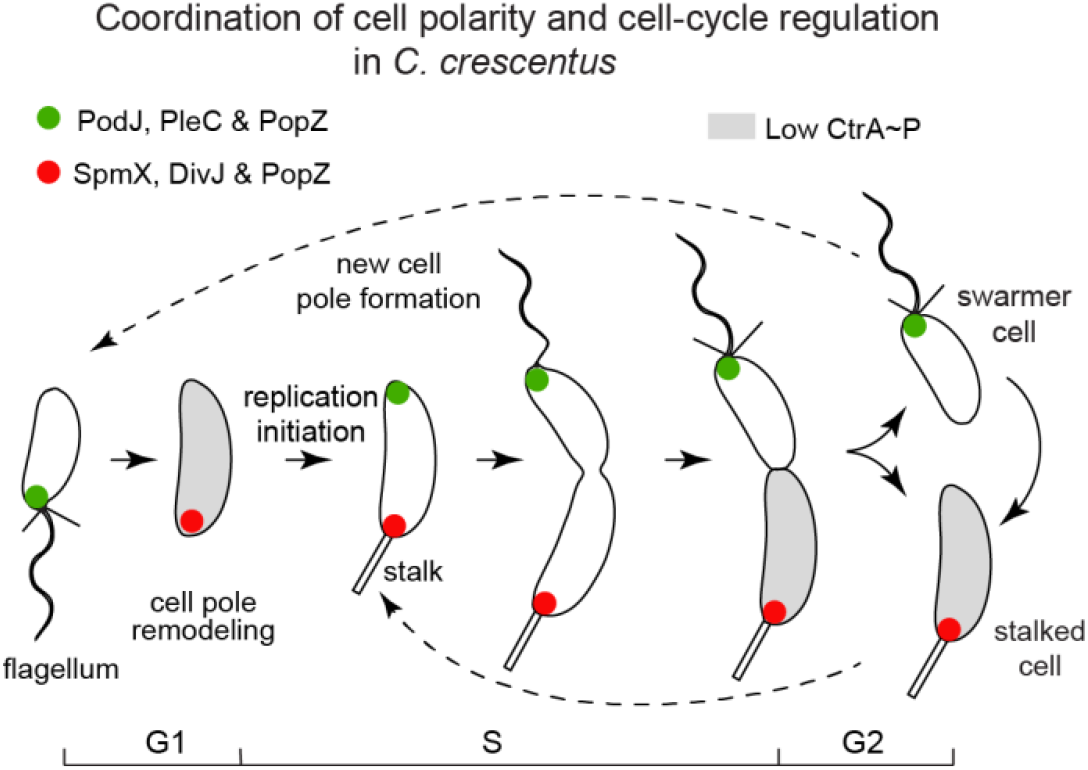
The PopZ and PodJ scaffold proteins are involved in the asymmetric accumulation of signaling proteins at the new cell pole. Swarmer cells of *Caulobacter crescentus* differentiate into stalked cells, which is associated with cell pole remodeling of a PodJ-rich signaling hub (green) into a SpmX-rich signaling hub (red). At the new pole of the stalked cells, a PodJ-rich signaling hub with scaffolding protein PopZ accumulates gradually upon initiation of replication. Cell division results in daughter cells that involved unequal inheritance of a PodJ-rich signaling hub in swarmer cell and a SpmX-rich signaling hub in stalked cell.

Amongst the client proteins asymmetrically polarized are a set of two-component signaling systems that collectively regulate the master regulator CtrA^3,6-10^. This intricate subcellular organization of CtrA regulators leads to selective CtrA phosphorylation at the new swarmer pole and dephosphorylation CtrA at the old stalked cell pole (Figure 1)^6,11^. Consequently, not only temporal^12^ but also spatial^13^ regulation of CtrA phosphorylation coordinate transcription of more than 90 developmental genes^14^. A scaffolding factor that is required for cell polarity is the protein PopZ. PopZ self-assembles as a micron-sized biomolecular condensate at each cell pole^13,15,16^. Single-molecule tracking experiments^13^, FLIP studies^16^, and *E. coli* reconstitution strategies^2,16,17^ have shown that PopZ dynamically recruits multiple distinct protein clients at each cell poles in pre-divisional cells^18^. However, the mechanisms that enable a common scaffold to promote the formation of two compositionally distinct biomolecular condensates remains unclear.

The new and old cell pole signaling hubs share some common clients, while others are selectively recruited to each signaling hub. The PopZ scaffold promotes bipolar accumulation of the histidine kinase CckA and its modulator DivL^16^. PopZ also serves as an attachment site for the ParB-*parS* centromere during chromosome segregation^15,18^. On the other hand, the histidine kinase DivJ specifically resides at the old cell pole, and the scaffolding protein SpmX mediates this specific recruitment. SpmX bridges the interaction between PopZ and DivJ, and can even nucleate the formation of new PopZ microdomains at ectopic poles upon overexpression^2^.

At the new cell pole, the scaffold proteins PopZ and PodJ play roles in polar assembly. Deletion of the PodJ scaffold results in failure to recruit PleC histidine kinase to the new cell pole^19,20^ and less monopolar accumulation of DivL at the new cell pole^21^. Moreover, Δ*podJ* strains exhibited moderate loss of the localization of PopZ’s client proteins at the new cell pole^21^. Downstream, this resulted in the down-regulation of the CtrA signaling pathway^21,22^ and reduced levels of the CtrA-regulated gene PilA^19,21,22^. Therefore, these previous studies suggest that similar to that of PopZ and SpmX at the old cell pole^2^, there are functional interactions between the PopZ and PodJ scaffolds at the opposite cell pole. Here we characterize the physical interactions between PopZ and PodJ within the new cell pole microdomain, and we demonstrate that PodJ-PopZ interaction coordinates the signaling transductions between their respective clients to ensure reliable asymmetric cell division.

## Results

### Newly translated PopZ accumulates at the new cell pole

A critical step in *C. crescentus* cell-cycle progression is the transition of PopZ from being localized exclusively at the old cell pole to accumulate at both cell poles. Given that PopZ scaffolds multiple cell-cycle factors^16,23^, we asked how the cell-pole condensates remain distinct during this change in localization patterns. One possible model is that PopZ can unbind its scaffold clients at the old cell pole and self-assemble as a separate matrix at the new cell pole. Alternatively, the accumulation of PopZ at the new cell pole may originate from the newly translated PopZ. In support of this second model, an increase in PopZ expression is observed at the same time as it is found that PopZ accumulates at the new cell pole^24^. We approached this question with a tandem fluorescent timer by fusing PopZ to one fluorescent protein that matures rapidly (sfGFP) and one that matures substantially more slowly (mCherry) (Figure 2A)^25^. Protein that exhibits high sfGFP fluorescence and weak mCherry fluorescence represents a newly translated protein. Protein that exhibits high sfGFP and high mCherry represents older protein. In past work applying this fluorescent timer approach, we demonstrated that in newborn swarmer cells, newly translated SpmX-mCherry-sfGFP accumulates at the old cell pole and ages as cells mature into pre-divisional cells^26^.

**Figure 2:**
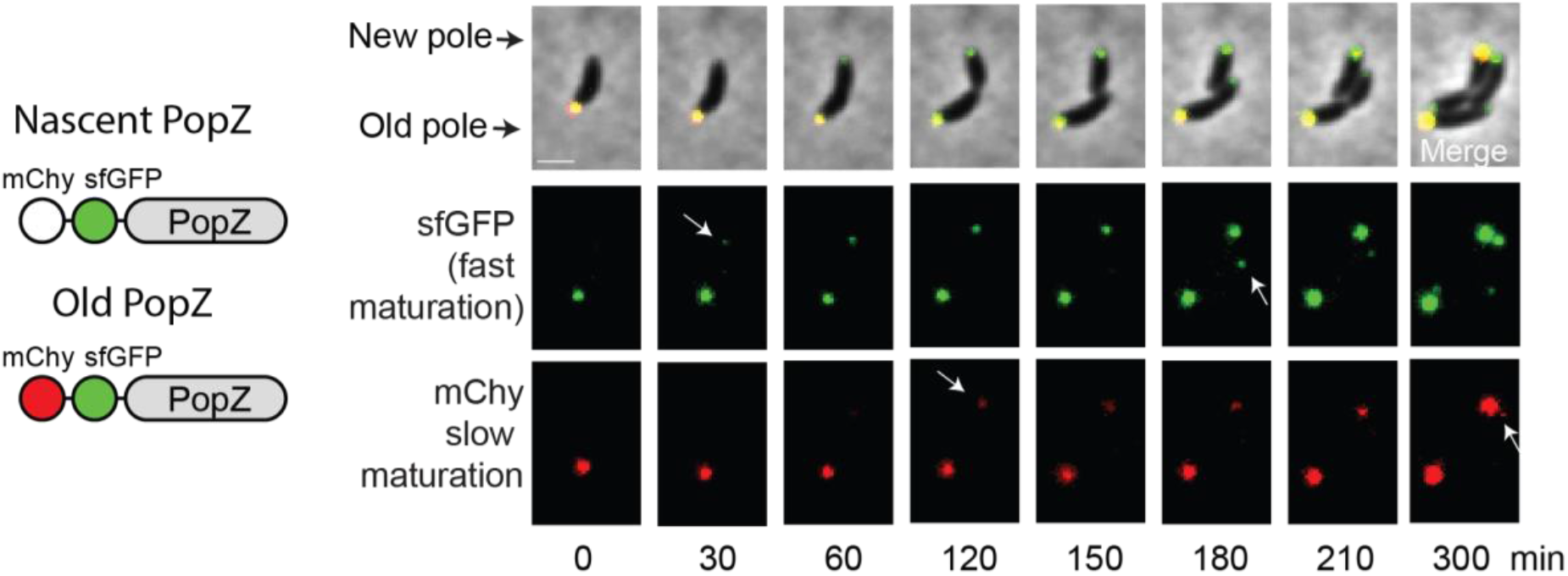
Newly translated PopZ localizes to the new cell pole in developing cells. mCherry-sfGFP-PopZ is expressed under the xylose promoter in NA1000 cells. mCherry (t_50_ maturation time of 45 min at 32°C) and sfGFP (t_50_ maturation time of 19 minutes at 32°C) chromophores mature at different times so newly synthesized PopZ will appear green and older synthesized PopZ appears as yellow. At time 0 min, the old pole shows both green and red indicating it is older yellow PopZ. At times 30-60 min a green PopZ focus appears at the opposite pole. At time 120 min the new foci contain both green and red fluorescence, indicating the subsequent maturation of the mCherry chromophore. Subsequently, in the second round of cell division, a green PopZ focus appears at the new cell pole of the divided cell at time 180 min as the newly translated PopZ appears at the new cell pole.

Time-course imaging on a synchronized *C. crescentus* population of mCherry-sfGFP-PopZ revealed that the new cell pole PopZ exhibited high sfGFP but weak mCherry signals at 30-minutes post-synchrony. In contrast, the old cell pole contained PopZ protein displayed both high sfGFP and mCherry signals (Figure 2A). At later time points in the cell cycle, 120-minutes post-synchrony, both high levels of sfGFP and mCherry can be observed at the new cell pole. This experiment indicated that older mCherry-sfGFP-PopZ is sequestered at the old cell pole, while the new cell pole is populated with newly translated PopZ protein. It is reasonable to presume that the sequestration of the old-new PopZ scaffolds may play a role in preventing the homogenization of PopZ and its clients at the new and old cell pole. Since PopZ subcellular localization abides by DNA occlusion mechanism^27^, a key question that follows is what promotes the accumulation of the newly translated PopZ at the new cell pole.

### PodJ regulates the amount of PopZ localized at the new cell pole

Previous studies have shown that ZitP^28^, TipN^29^, and ParA^30^ play redundant roles in the accumulation of PopZ at the new cell pole but implicate one or more additional unknown players. We hypothesized that a PopZ-PodJ scaffold-scaffold interaction may occur since only PodJ could provide the recruitment capability in these players at the new cell pole^19-22,31^.

We observed that sfGFP-PodJ was able to accumulate at the poles in over 90% of cells in the Δ*popZ* strain (Figure S1A). However, we also observed an increase in cells exhibiting bipolar localization (Figure S1A). This increase in PodJ bipolar accumulation could be due to differences in PodJ protein levels or changes levels of PodJ proteolysis. For example, in strains lacking the PodJ protease PerP, the number of cells that exhibit bipolar accumulation of PodJ substantially increased (Figure S1B), consistent with past observations^32^. Notably, we did not observe an increase in diffuse PodJ in the Δ*popZ* strain. Therefore PodJ’s ability to accumulate at the cell poles is independent of the PopZ scaffold.

We did, however, observe a 3-fold reduction of PopZ accumulation at the new cell pole in the Δ*podJ* versus wild-type strain (Figure 3A). Expression of sfGFP-PodJ from the chromosomal xylose locus recovered the robust PopZ accumulation at the new cell pole (Figure 3A). These results suggest that PodJ plays a role in regulating the amount of PopZ accumulation at the new cell pole. We also observed that cells without full-length PodJ also showed a decrease in total cell mCherry-PopZ intensity (Figure 3C). This suggests that deleting the native *podJ* gene may alter PopZ transcription levels. Hence, the decreased mcherry-PopZ accumulation at the new cell pole may be due to reduced expression of mCherry-PopZ or loss of physical recruitment. We therefore, examined the distribution of PopZ in cells by constitutive expression of mCherry-PopZ from the vanillate locus. Also, a 4-fold reduction in the fraction of mCherry-PopZ signal at the new cell pole was observed in Δ*podJ* compared to the wild-type strain (Figure S2A, S2B). Therefore, higher levels of PopZ expression alone are not capable of rescuing the loss of PopZ accumulation at the new cell pole.

**Figure 3:**
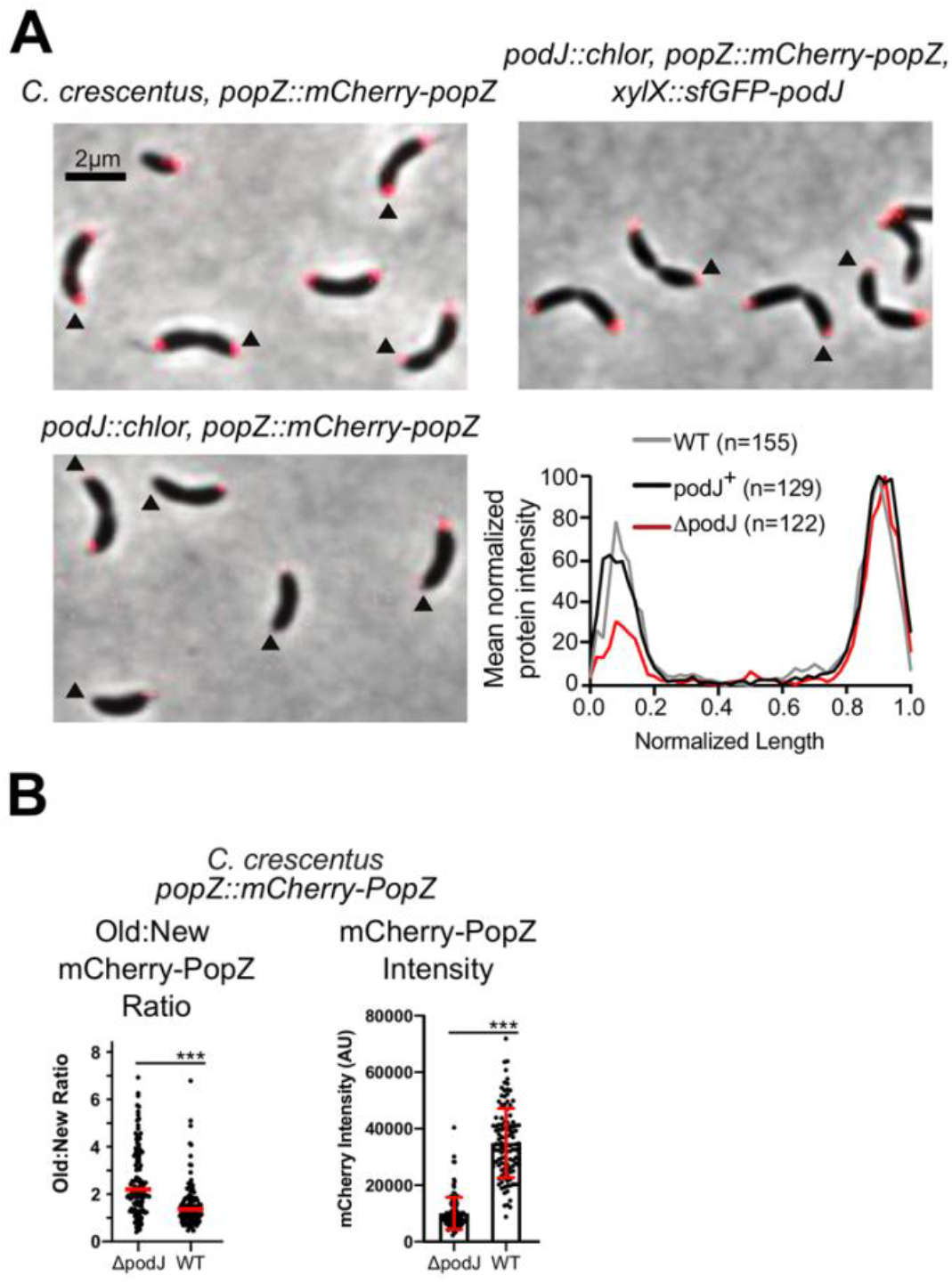
PodJ regulates PopZ assembly at the new cell pole. Analysis of the impact of the Δ*podJ* upon mCherry-PopZ’s localization pattern in *C. crescentus*. The expression of the sole copy *popZ* was induced from PopZ’s native promoter in the chromosome. (A) mCherry-PopZ localization in predivisional cells in the wild-type (bipolar) versus the *podJ* deletion *C. crescentus* (monopolar). The quantitative analysis reveals a substantial reduction of PopZ abundance at the new cell pole of Δ*podJ* predivisional cells. Bars, 2 μm. (B) Comparison of the percentage of cells displaying bipolar PopZ in wild-type and Δ*podJ*. Analysis of Old/New cell pole ratio and total cell intensity of mCherry-PopZ in different PodJ backgrounds. *** indicates *p* < 0.0001. Red line indicates mean. Red bars indicated mean ± standard deviation. Statistical analysis done using student’s t-test.

We also performed time-lapse microscopy experiments to examine the mCherry-PopZ localization throughout the cell cycle starting with a synchronized population of swarmer cells (Figure S2C). Images were acquired every minute, and kymographs were constructed to show the fluorescence intensity along the cell body over time. In wild-type cells, robust mCherry-PopZ foci accumulated at the new cell pole approximately 40 minutes post-synchrony (Figure. S2C, Movie S1). However, in a Δ*podJ* strain, we detected significantly reduced signal at the new cell pole (Figure S2C, Movie S2). Moreover, a subset of nascent swarmer cells that lacked any observable PopZ focus were observed (Figure S2D). This loss of PopZ could be complemented by expressing sfGFP-PodJ (Figure S2D). Amongst these swarmer cells, we found 91% of cells ultimately accumulated PopZ at the correct, old cell pole (Figure S2E). We observed that 9% of these cells accumulated PopZ at the new cell pole after inheriting no PopZ (Figure S2E). Thus, this subpopulation of swarmer cells exhibited an abnormal switching of the polarity axis.

This observed reduction in PopZ new cell pole accumulation mirrors loss other redundant factors (TipN^33^ and ZitP^28^) that play roles in promoting PopZ new cell pole accumulation. This redundancy in PopZ recruitment likely reflects how deletion of *podJ* does not result in phenotypes seen in cells with *popZ* deleted^15^. Collectively, these results suggest that the degree and the time of PopZ accumulation at the new cell pole depends on PodJ, but PodJ cell pole accumulation is independent of PopZ.

### PodJ deletion impacts ParB segregation in a subset of cells

Past work from Brun and co-workers have shown that the PopZ client CckA exhibits reduced new cell pole localization when *podJ* is deleted or truncated^21^. Another critical role of PopZ is to tether the ParB/origin segregation complex at the cell poles^15^. The robust tethering of ParB to the cell poles involves simultaneous interactions with numerous ParB/*parS* complexes^17,34^. Therefore, we investigated if the reduction of PopZ accumulation at the new cell pole impacted ParB tethering. Previously, Bowman and co-workers demonstrated that ParB was tethered more stably at the new cell pole than at the old cell pole after chromosome segregation^23^. We observed that ParB-CFP was able to readily accumulate at the new cell pole, while ParB-CFP foci were more mobile at both the swarmer and stalk pole, with the greater change in mobility at the swarmer cell pole when cells lacking PodJ (Figure 4A). This observation suggests that a PodJ mediated recruitment of PopZ impacts the dynamics of the ParB/origin complex at the cell pole This close association of ParB with the cell poles is likely due to the lower degree of subcellular accumulation of PopZ at the new cell pole. Alternatively, it may also suggest that the Pod-PopZ interaction allosterically impacts the PopZ-ParB interaction.

**Figure 4:**
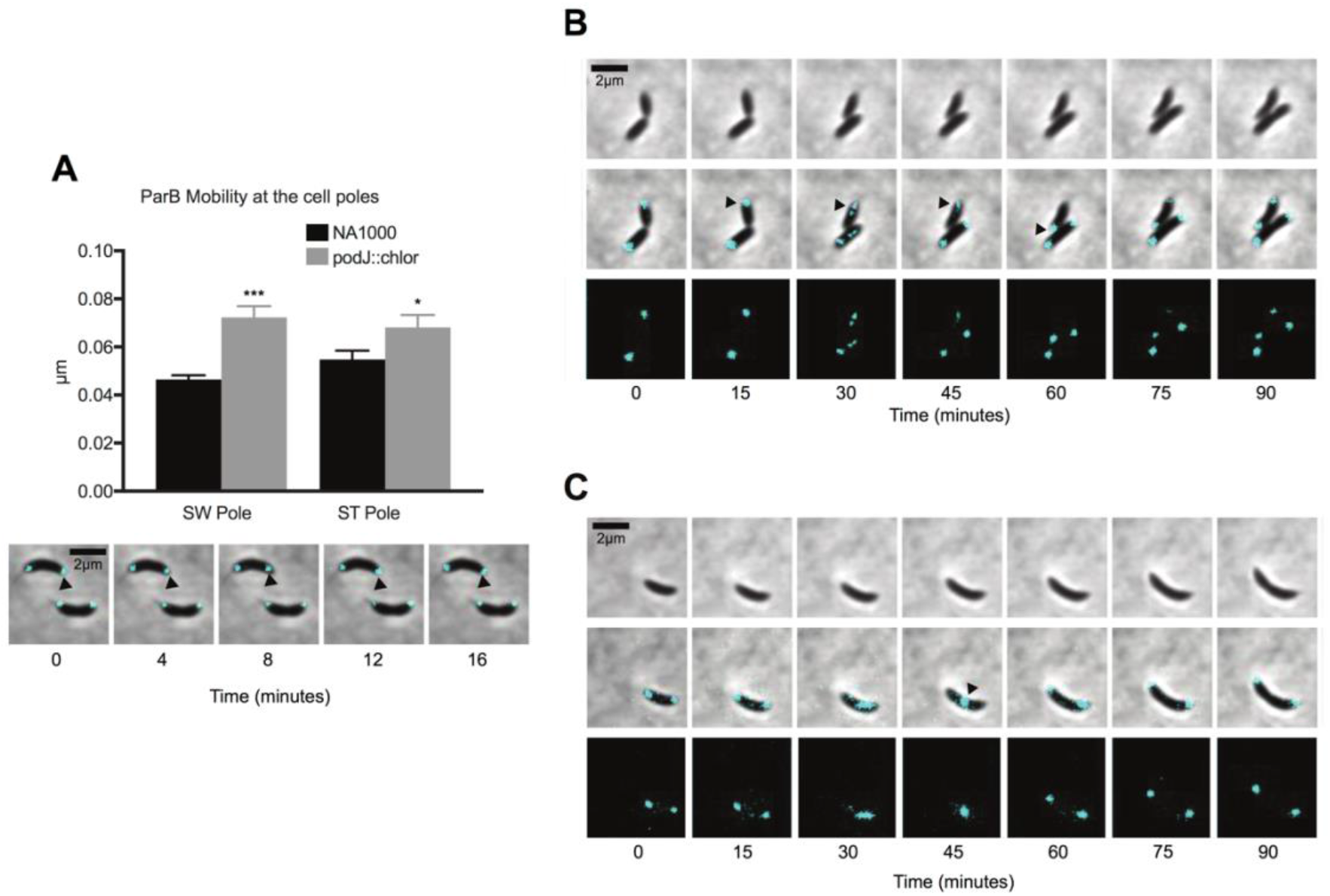
*C. crescentus* strains lacking PodJ exhibit chromosome segregation defects. (A) Analysis of ParB foci mobility at the cell poles in wild-type versus Δ*podJ* strains. Cells shown are Δ*podJ* background. *** indicates *p* < 0.0001 and * indicates *p* < 0.05. Student’s t-test used for statistical significance. (B and C) Observed chromosome translocation defects in the Δ*podJ* strain.

Additionally, we observed that 35% of cells displayed ParB focus detachment phenotypes in the *podJ* deletion strain at both cell poles. In the most prevalent cases, the ParB focus would translocate across the cell to the new cell pole before chromosome duplication (Figure 4B). This premature centromere translocation results in the reversal of the inherited cell polarity axis. In another case, we observed new and old cell pole ParB foci coalescing into a single focus at the middle cell, then separating back to the cell poles (Figure 4C). Consistent with the mobility analysis results (Figure 4A), these phenotypes suggest the PodJ recruitment of PopZ facilitates robust PopZ-ParB chromosome tethering at the new cell pole.

Given that ParB also directly interacts with the cell division inhibitor protein MipZ^35^, we examined the impact of the *podJ* deletion upon MipZ and FtsZ. These ParB segregation defects also resulted in a less robust MipZ localization at the cell poles and a more diffuse FtsZ Z-ring assembly (Figure S3A, S3B). Overall in the *podJ* deletion strain, cells were viable as chromosome segregation, and division processes remained mostly functional. However, PodJ’s interaction with PopZ seems to fine-tune chromosome segregation such that it avoids polarity axis inversions.

### PodJ promotes bipolarization of PopZ in E. coli

To determine if PodJ and PopZ interact directly, we heterologously co-expressed PopZ and PodJ scaffolds in *E. coli (*Figure 5A, 5B). Notably, the γ-proteobacterium *E. coli* is highly divergent from the alphaproteobacterium *C. crescentus* and does not contain any *C. crescentus* polarity protein homologs. *E. coli* has thus been used extensively as an orthologous system for testing *C. crescentus* protein-protein interactions^15,16,27,28^. A previous screen of PopZ interaction partners indicates that PopZ and PodJ were only partially co-localized when co-expressed in *E. coli*^16^ despite their co-localization in *C. crescentus*. This previous study utilized a C-terminal fluorescent protein fusion to PodJ, while previous PodJ studies have used an N-terminal fluorescent protein fusion of PodJ^32,36^. Therefore, we hypothesized that the C-terminal fluorescent protein fusion might impact PodJ localization and therefore disturb PodJ-PopZ binding. To test this idea, we heterologously expressed an N-terminal fluorescent fusion protein of PodJ in *E. coli*. As shown in Figure 5A, YFP-PodJ exhibited readily bipolar localization in about 80% of *E. coli* cells (Figure 5A, S4). PopZ accumulates at a single cell pole in about 75% of cells when expressed alone, as observed in past studies^18,27^ (Figure 5A, S4). However, mCherry-PopZ co-localized in a bipolar pattern when co-expressed with YFP-PodJ (Figure 5A, 5B). Therefore, these experiments indicated that PodJ could bipolarize PopZ in *E. coli* (Figure 5, S4). Interestingly, this PodJ-mediated bipolarization of PopZ might be a general feature of membrane-bound PopZ client proteins as SpmX^2^, ZitP^28^, and DivL^16^ all can bipolarize PopZ in *E. coli*.

**Figure 5:**
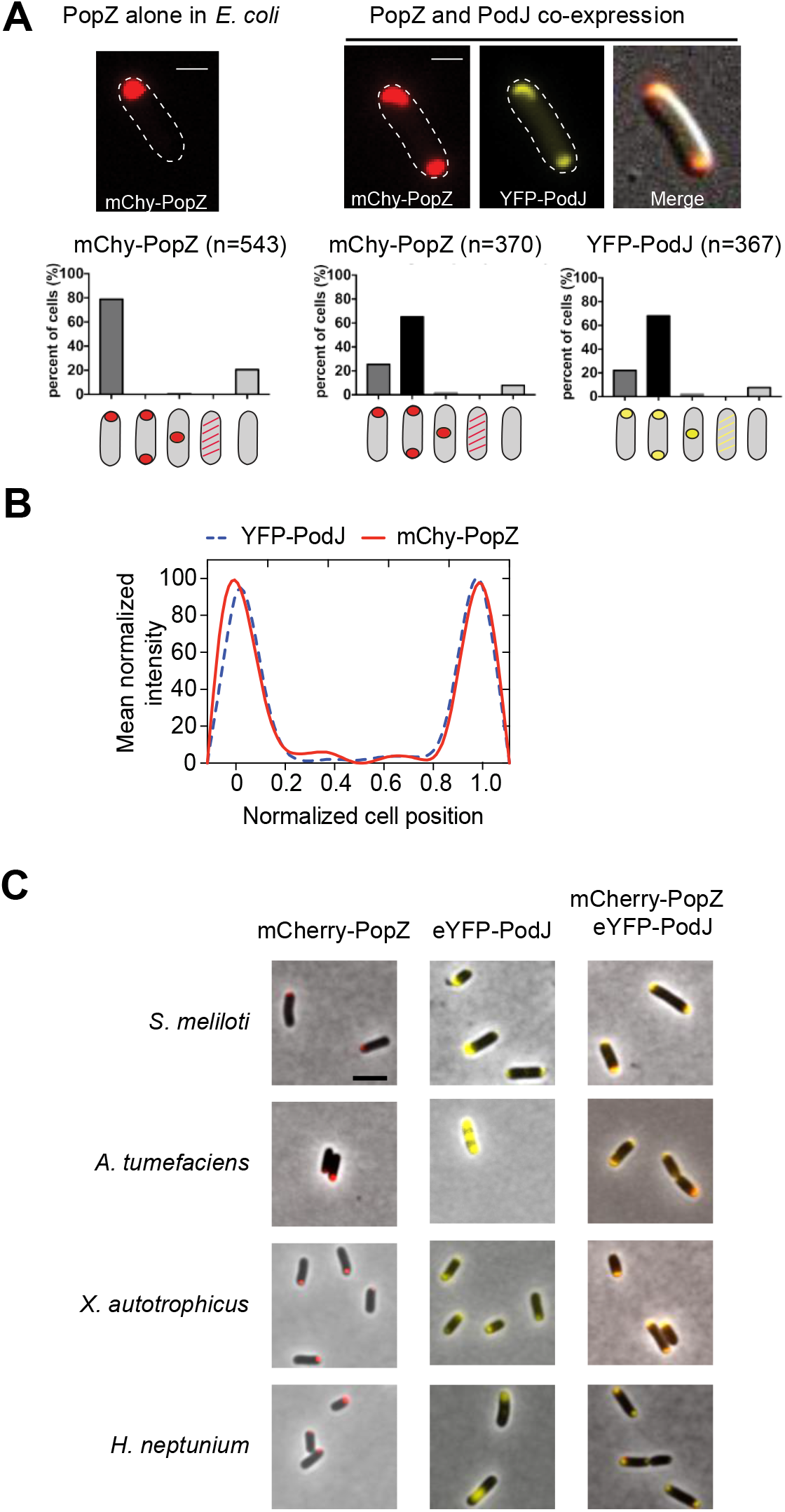
PodJ bipolarizes PopZ when expressed in *E. coli*, via an interaction conserved across alphaproteobacteria. **(A)** Heterologous expression of YFP-PodJ and mCherry-PopZ in *E. coli*. Co-expression with PodJ causes bipolar PopZ accumulation in *E. coli*. (B) Mean protein intensity of YFP-PodJ and mCherry-PopZ versus cell length (n = 370). The signal intensity was normalized with the highest value as 100% in each strain. **(C)** Co-expression of PopZ-PodJ scaffold pairs from *Sinhorhizobium meliloti, Agrobacterium tumefaciens, Xanthobacter autotrophicus*, and *Hyphomonas neptunium*. All PopZ homologs accumulate specifically at one cell pole when expressed alone. Co-expression of PopZ together with PodJ results in co-localized PopZ-PodJ bipolar localization.

### PopZ-PodJ interaction is conserved amongst alphaproteobacteria

A subset of alphaproteobacteria encodes both PopZ and PodJ scaffolding proteins. Notably, in the alphaproteobacteria *Agrobacterium tumefaciens*, past studies have demonstrated a strong genetic interaction between PodJ and PopZ^37,38^. However, from these prior studies, it remains unclear if *At*PodJ and *At*PopZ interact directly or indirectly. To test this idea, we expressed PodJ fusion proteins from select alphaproteobacteria together with their corresponding PopZ variants in *E. coli* (Figure 5C). Each mCherry-PopZ homolog accumulated at a single cell pole when expressed alone, similar to *Cc*PopZ (Figure 5C). Each YFP-PodJ variant accumulated at the cell poles, but compared to *Cc*PodJ, the variants displayed heterogeneity in their subcellular localization pattern. However, in each case, we observed that co-expression with PodJ results in bipolarization of PopZ (Figure 5C). These experiments indicate that the interaction between PopZ and PodJ is direct and conserved amongst alphaproteobacteria that contain both PopZ and PodJ.

### PopZ interacts directly with PodJ’s CC4-6 domain

To determine the PopZ binding site within PodJ, we screened the capability of PopZ to bind to the library of PodJ domain deletion variants through co-expression in *E. coli* (Figure 6A, S4). We considered the following outcomes as an indication of a PopZ interaction with the PodJ variants: (1) the two proteins are 100% co-localized, and (2) the localization pattern of either protein is changed after co-expression. We found that the deletion of the C-terminal periplasmic domain or the intrinsically disordered PSE domain in PodJ did not disrupt the PodJ-PopZ interaction (Figure 6A, Figure S4). In contrast, the deletion of the CC4-6 domain disrupted PopZ co-localization with PodJ (Figure 6A). We then expressed YFP-CC4-6 alone and observed that it was diffuse through the cytoplasm in *E. coli*. However, it co-localized with mCherry-PopZ at the cell pole when co-expressed in *E. coli* (Figure 6A). These data indicate that coiled-coil 4-6 in PodJ is critical for co-localization with PopZ in *E. coli*.

**Figure 6:**
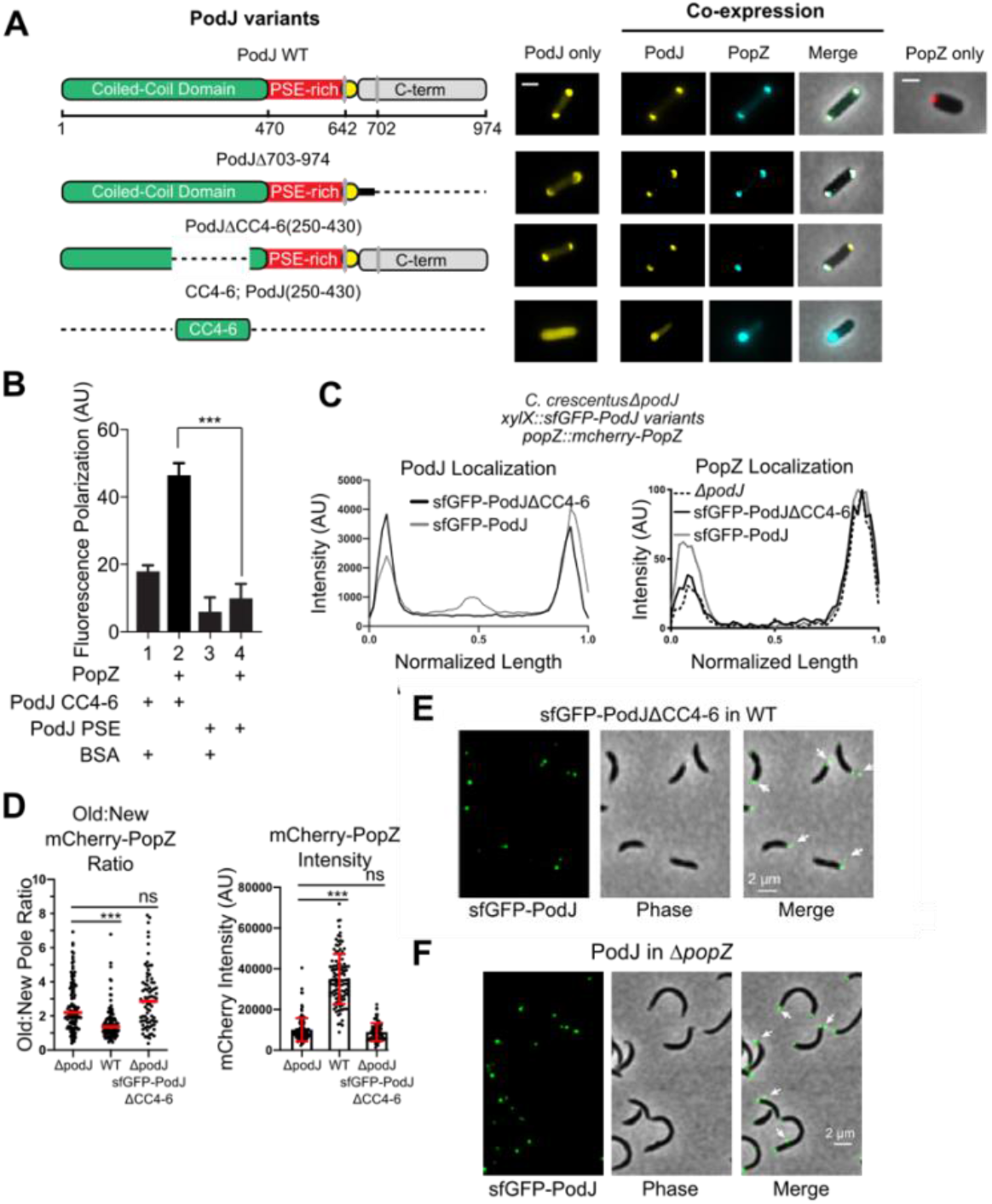
PopZ binds directly to the coiled-coil 4-6 region of PodJ. **(A)** Co-expression of PodJ variants together with PopZ in *E. coli* reveals that the coiled-coil 4-6 region in PodJ is necessary for the interaction with PopZ (please refer to Figure S4 for more details). (B) Fluorescence polarization binding assay of the BODIPY dye-labeled PodJ_PSE or PodJΔCC4-6 mixed with 10 µM PopZ, using BSA as a negative control. PopZ binds specifically to the CC4-6 domain of PodJ. However, PopZ does not bind to its PSE-rich domain. (C) Fluorescent plots normalized by cell length where 0.0 is the new cell pole, 1.0 is the old cell pole with the expression of sfGFP-PodJ variants from the xylose promoter in *C. crescentus*. These *ΔpodJ* cells are also expressing mCherry-PopZ from the *popZ* promoter. (D) Analysis of Old/New cell pole ratio and total cell intensity of mCherry-PopZ in different PodJ backgrounds. *** indicates *p* < 0.0001. Red line indicates mean. Red bars indicated mean ± standard deviation. Statistical analysis done using student’s t-test. (E) Loss of PodJ-PopZ interaction results in stalk-pole specific foci that contain PodJΔCC4-6 protein. (F) sfGFP-PodJ in *ΔpopZ* cells. Arrows indicate sfGFP-PodJ found outside of the cell or in non-polar regions of the cell.

To confirm that this PopZ-PodJ protein-protein interaction is direct, we performed *in vitro* fluorescence polarization assays to detect PopZ-PodJ binding. In these assays, we mixed 16 µM PopZ together with 100 nM BODIPY-PodJ CC4-6 or BODIPY-PodJ PSE fluorescently labeled proteins. As shown in Figure 6B, PopZ bound to PodJ CC4-6 but did not bind to the PodJ PSE construct. Both the *E. coli* heterologous expression assays and *in vitro* biochemical assays show that the coiled-coil 4-6 region of PodJ is the site of interaction with PopZ.

### PodJ-PopZ interaction regulates PopZ new pole localization and loss of PodJ from cells

In *C. crescentus* Δ*podJ*, we observed that the expression of sfGFP-PodJΔCC4-6 was able to localize at the new cell pole (Figure 6C). One notable difference is that sfGFP-PodJΔCC4-6 exhibited an increased mid-cell accumulation versus sfGFP-PodJ. A second critical difference is that sfGFP-PodJΔCC4-6 recruited about 2-fold less PopZ to the new cell pole than the expression of sfGFP-PodJ (Figure 6C, 6D). A comparison of PopZ cell pole intensity ratio (old/new) in the wild-type strain versus the PodJΔCC4-6 strain and the *ΔpodJ* strain shows the ratio increases in cells lacking PodJ with a functional PopZ binding site (Figure 6D). Taken together, these results suggest that the PodJ CC4-6 binding site contributes to PopZ accumulation at the new cell pole.

To our surprise, we observed that sfGFP-PodJΔCC4-6 foci outside of the cell, specifically at the old cell pole (Figure 6E). We also observed a similar phenomenon when expressing sfGFP-PodJ in *popZ* deletion strain (Figure 6F). One possible explanation is the formation of minicells, which have been described in previous studies of PopZ^27^ and SpmX^39^ mutant strains, and SpmX overproducing cells^2^. Previous work from Thanbichler et al. demonstrated that mini-cell formation is commonly the result of chromosome detachment errors, as observed in MipZ mutant strains^35^. This is partially consistent with our observation of increased ParB mobility at the cell poles and abnormal ParB translocation events (Figure 4B, 4C). However, given the role of the PopZ-PodJ interaction at the cell poles, we would expect mini-cell formation to occur equally at both poles especially at the new cell pole. Ebersbach et al. previously showed that minicells produced in the *popZ* deletion strain occur exclusively at the new cell pole^27^. In contrast, in the *popZ* deletion strain we observed extracellular PodJ-rich foci exclusively at the old cell pole (Figure 6D). In addition, these foci were significantly smaller than mini-cells and not observable by phase in most cases. Another possibility for the observed extracellular PodJ is that PodJ or a complex, including PodJ, is secreted from the cell body. This could occur via the CpaC outer membrane secretion channel, which remains assembled at the old cell pole after facilitating the secretion of the PilA pilin protein at the new cell pole early in the cell cycle ^19,40^. Notably, a second factor that plays a role in pilus assembly, CpaE, is recruited to the cell pole by the PodJ scaffolding protein and is required for CpaC localization^19,40^. Investigation of this process and its relevance to cell-cycle regulation will require further genetic studies. Regardless of the mechanism of PodJ loss, these results suggest that PopZ-PodJ interaction is critical for robust tethering of the chromosome at the cell poles (Figure 4) and prevention of loss of PodJ from the cell body (Figure 6).

## Discussion

Recently, biomolecular condensation has emerged as an organizing principle of the bacterial cytoplasm^13,41-44^. Moreover, it has been shown that the scaffolding protein PopZ play an essential role in the formation of two biomolecular condensates at each cell pole^13,16^. Here we have discovered a direct and conserved interaction between the PopZ and PodJ scaffolds (Figure 6B, S5) influences the composition and the size of biomolecular condensates at the new cell pole (Figure 3, S2)^13^. In the absence of PodJ, we observed a 3 to 4-fold reduction in the amount of PopZ that localized to the new cell pole (Figure 3, S5). This reduction in new cell pole localized PopZ also had an impact upon tethering of ParB to the cell poles. We observed erroneous ParB translocations from the old cell pole to the new cell pole before chromosome duplication in the *podJ* deletion strain (Figure 4B, S5). Therefore, PodJ plays a role in ensuring cells inherit and maintain their polarity axis. Overall, the observed segregation and division phenotypes were mild, indicating that PopZ has the ability to self-assemble at the new cell pole as other redundant proteins play a role in PopZ new-pole promotion (Figure S5)^28,30^.

A key event in *C. crescentus* asymmetric division is the formation of a signaling hub at the new cell pole that is compositionally distinct from the old cell pole (Figure S5). Previous fluorescence recovery after photobleaching (FRAP) experiments^13,16^ and single-molecule tracking experiments^15^ collectively indicate that PopZ is sequestered at the old poles for long periods of time. From these past experiments, we hypothesized that PopZ accumulation at the new cell pole primarily occurs through the assembly of newly translated PopZ. To distinguish newly translated from older PopZ, we applied a fluorescent-timer approach. These fluorescent-timer protein fusions demonstrated that newly translated protein was enriched at the new cell pole (Figure 2), while old PopZ protein was sequestered mainly at the old cell pole. Thus the combination of single-molecule tracking (< 1 min)^15^, FRAP (0-10 min)^13,16^, and fluorescent timer data (>10 min) (Figure 1) allow tracking of protein over a range of timescales, and each of these methods suggests that sequestration of static PopZ assemblies play a role in preventing the scrambling of contents at the cell poles.

Super-resolution imaging of the cell poles suggests that the molecular organization is well mixed at the spatial resolution of approximately 20 nm^45^. In the absence of protein-protein interaction information, the PopZ-CckA-DivL and PodJ-PleC complexes could either be interacting and well mixed or non-interacting and phase-separated into discrete clusters at the new cell pole. Our observation of a direct-scaffold interaction between PodJ and PopZ (Figure 3, 6, S2) likely mediates placement of PleC, CckA, DivL as a well-defined signaling complex in alphaproteobacteria (Figure 5). This proximity would support previously proposed models in which PleC’s dephosphorylation of DivK∼P may generate localized zones of unphosphorylated DivK∼P^11,19^. In contrast, simple co-localization of signaling proteins at the cell poles as heterogeneous clusters and without direct interactions may not overcome the rapid DivK diffusion rates that generate shallow DivK∼P gradients across the cell^46^.

More broadly, recent work has identified an array of scaffolds that promote the organization of bacterial cytoplasm from signaling biochemistry^16,45^ to RNA biochemistry^41^ through self-assembly as biomolecular condensates. Key questions remain as to the factors that promote co-assembly, phase separation, and compositional control of these bacterial biomolecular condensates. Future studies will be needed to determine if PodJ can self-assemble and whether it is homogenously integrated at the membrane-PopZ microdomain surface. In contrast, the absence of these scaffold-scaffold interactions, and other yet to be learned mechanisms, may facilitate phase separation of distinct biomolecular condensates. For example, C. *crescentus* contains three known spatially resolved biomolecular condensates: BR-bodies involved in mRNA decay dispersed in the cell-body^41^, and two PopZ-mediated assemblies at opposite cell poles^16^. System-level understanding of the bacterial cytoplasm organization within these biomolecular condensates will center on understanding the breadth of scaffold-scaffold interactions.

## Acknowledgments

We thank Jared Schrader and Tom Mann for providing critical reviews of the manuscript. We also thank Lucy Shapiro for providing critical *C. crescentus* strains that supported this study.

## Methods

### Bacterial Strains

All experiments were performed using *Caulobacter crescentus* NA1000 (also known as CB15N) and *Escherichia coli* BL21. *E. coli* BL21 was purchased from Promega. *C. crescentus* NA1000 was a kind gift from Dr. Lucy Shapiro (Stanford University School of Medicine). More strains and expression plasmids used in this study are listed in Table S1. All relevant primers are given in detail in Table S2. Plasmid and strain construction are described in the supplemental information. Transformations and phage transductions were carried out as described^47^.

### Growth Conditions and Inducer Concentrations

*C. crescentus* strains were grown at 28°C in PYE (peptone yeast extract) or M2G (minimal medium supplemented with glucose)^47^. When needed, *C. crescentus* cells were synchronized as described^48^, and swarmer cells were harvested by Percoll density-gradient centrifugation. *E. coli* strains used for protein purifications and microscopy experiments were grown at 37 °C in LB medium unless otherwise stated. When required, protein expression was induced by adding 0.002-0.5 mM Isopropyl β-D-1-thiogalactopyranoside (IPTG) or 0.5-10 mM arabinose in *E. coli*, and 0.003%–0.3% xylose or 0.05-0.5 mM vanillic acid in *C. crescentus* unless otherwise stated. The induction time for microscopy experiments is 2 hours in *E. coli* and 3 hours in *C. crescentus*. Generalized CR30 phage transduction was performed as described^47^.

### Phase Contrast, DIC, and Epifluorescence Microscopy

Cells were imaged after being immobilized on a 1.5% agarose pad containing corresponding inducers when required. Phase microscopy was performed by using a Nikon Eclipse T*i*-E inverted microscope equipped with an Andor Ixon Ultra DU897 EMCCD camera and a Nikon CFI Plan-Apochromat 100X/1.45 Oil objective. DIC (differential interference contrast) microscopy was performed using the same microscope and camera but with a Nikon CFI Plan-Apochromat 100X/1.45 Oil DIC objective with a Nikon DIC polarizer and slider in place. The excitation source was a Lumencor SpectraX light engine. Chroma filter cube CFP/YFP/MCHRY MTD TI was used to image ECFP (465/25M), EYFP (545/30M), and mCherry (630/60M). Chroma filter cube DAPI/GFP/TRITC was used to image EGFP, sfGFP, and mNeonGreen (515/30M). Images were collected and processed with Nikon NIS-Elements AR software.

### Time-lapse Microscopy

sfGFP-PodJ, mCherry-PopZ, or SpmX-mCherry were tracked using phase and fluorescence microscopy. During time-lapse experiments, phase and fluorescence images were taken in 1 min intervals for sfGFP-PodJ, mCherry-PopZ, and SpmX-mCherry for 1-2 cell divisions (∼ 4 h). ParB-CFP fast time-lapses images were recorded every 4 minutes over 20 minutes. Long ParB-CFP time-lapses were recorded every 15 minutes for 3-4 hours. The imaging system used was the Nikon Eclipse T*i*-E microscope equipped with an Andor Ixon Ultra DU897 EMCCD camera and NIS-Elements software. *C. crescentus* cells with corresponding expression gene were grown to the early-log phase in M2G or PYE medium (OD_600_ = 0.2), and then induced by xylose or vanillic acid for 2 hours before synchronization. Swarmer cells were isolated from the culture by centrifugation (20 mins at 11,000 rpm, 4°C) after mixture with 1 volume of Percoll (GE Healthcare). The synchronized swarmer cells were pipetted onto an agarose (2%) pad containing medium with inducers and sealed with wax. NIS-Elements software was used to align time-lapse images post-acquisition.

### ParB-CFP tracking analysis

MicrobeJ^49^ was used to track ParB-CFP foci during fast time-lapse experiments. Predivisional cells that had already segregated a ParB-CFP focus to the new cell pole were at t=0 were analyzed. Maxima were tracked, and the raw distance changes for each 4-minute difference were averaged for new and old cell pole ParB-CFP foci. Averages for two separate experiments were pooled and plotted. A student’s t-test was used to determine statistical significance.

### Fluorescence Intensity Profile Analysis

sfGFP-PodJ variants expressing mCherry-PopZ from the native PopZ promoter were imaged using the above methods. After imaging, predivisional cells expressing sfGFP-PodJ variants were oriented by visualization of the stalk. The average fluorescence intensity profile using normalized cell length was generated using MicrobeJ^49^ with the new pole at 0.0 and old pole at 1.0. mCherry-PopZ was made in the same way in the same strains. MipZ and FtsZ analysis were performed in the same way.

### Purification of PodJ and PopZ

Protein expression of all PodJ variants followed the same protocol and is described in detail below for PodJ (1-635). To purify the cytoplasmic portion of PodJ(1-635), Rosetta (DE3) containing plasmid pwz091 was grown in 6 liters LB medium (20 µg/ml chloramphenicol and 100 µg/ml ampicillin) at 37°C. The culture was then induced at an OD600 of 0.4–0.6 with 0.5 mM IPTG overnight at 18°C. The cells were harvested, resuspended in the lysis buffer (50 mM Tris-HCl, 700 mM KCl, 20 mM Imidazole, 0.05% dextran sulfate, pH 8.0), in the presence of protease inhibitor cocktail tablets without EDTA (Roche).

The cell suspension was lysed with three passes through an EmulsiFlex-C5 cell disruptor (AVESTIN, Inc., Ottawa, Canada), and the supernatant was collected by centrifuging at 13000 *g* for 30 min at 4°C. Also, the insoluble cell debris was resuspended by the recovery buffer (50 mM Tris-HCl, 1000 mM KCl, 20 mM Imidazole, 0.05% dextran sulfate, pH 8.0) and its supernatant was collected as well as the previous centrifugation. The combined supernatants were loaded onto a 5 ml HisTrap™ HP column (GE Healthcare) and purified with the ÄKTA™ FPLC System. After washing with 10 volumes of wash buffer (50 mM Tris-HCl, 300 mM KCl, and 25 mM imidazole, pH 8.0), the protein was collected by elution from the system with elution buffer (50 mM Tris-HCl, 300 mM KCl, and 500 mM imidazole, pH 8.0), and concentrated to a 3 ml volume using Amicon Centrifugal Filter Units, resulting in > 95% purity. All PodJ variants were dialyzed with a buffer containing 50 mM Tris-HCl (pH 8.0), 300 mM KCl, and then aliquoted to a small volume (100 µl) and kept frozen at −80°C until use.

His-PopZ was expressed and purified the same as described ^17^.

### Fluorescence Polarization Assay

To label PodJ_PSE (471-635) and PodJ_CC4-6 (250-430), we cloned a cysteine just after the 6X-His-tag proteins at the N-terminal of each protein. PodJ_PSE (Cys) and PodJ_CC4-6 (Cys) expression and purification followed the same protocol as PodJ mentioned above. These two proteins were labeled at the cysteine using thiol-reactive BODIPY™ FL N-(2-Aminoethyl) Maleimide (Thermo Fisher). The proteins were mixed with 10-fold excess BODIPY™ FL N-(2-Aminoethyl) Maleimide and allowed to react for 2 hours at room temperature, and the unreacted dye was quenched with mercaptoethanol (5% final concentration). The labeled proteins were purified via dialysis to remove unreacted fluorescent dye (5 times, 500 ml buffer, and 30 mins each).

Fluorescence polarization binding assays were performed by mixing 100 nM labeled proteins with 0, 0.25, 0.5, 1, 2, 4, 8, 16 µM partner protein (PopZ or BSA) for 45 minutes to reach binding equilibrium at the room temperature. Fluorescent proteins were excited at 470 nm, and emission polarization was measured at 530 nm in a UV-vis Evol 600 spectrophotometer (Thermo). Fluorescent polarization measurements were performed in triplicates, and three independent trials were averaged with error bars representing the standard deviation.

### Quantification and Statistical Analyses

FIJI/ImageJ^50, 51^, and MicrobeJ ^49^ were used for image analysis. The number of replicates and the number of cells analyzed per replicate is specified in corresponding legends. All experiments were replicated at least 2 times, and statistical comparisons were carried out using GraphPad Prism with two-tailed Student’s t-tests. Differences were considered to be significant when *p* values were below 0.05. In all figures, measurements are shown as mean ± standard deviations (s.d.).

### Kymograph Analyses

Kymographs of fluorescence intensity was obtained by using the built-in kymograph function of MicrobeJ^49^. The background signal was subtracted before the kymograph analysis, and the observation of stalk at the pole of *C. crescentus* cell was defined as the old pole. The predivisional cell was selected as the start point in Figure 1C and Figure 3C. In Figure 1C, another round of kymograph analysis was performed after the first cell division. The new pole **b** became the old pole after cell division and another two new poles (**c** and **d**) were formed.

### Calculation of Subcellular Co-Localization with PodJ variants

To interpret the co-localization ratio in Figure 4C and Figure S2, we used strict criteria to calculate how the proteins interact with the PodJ variants, *i*.*e*., (I), the localization patterns of the interaction proteins are changed after co-expression. (II), the two proteins are 100% co-localized at the pole (binding) or drive each other apart from the pole (dispersion). Failure to meet either of these two criteria means the interaction of the two proteins is undetermined. About 200 cells were analyzed for each interaction set.

## References

1 Good, M. C., Zalatan, J. G. & Lim, W. A. Scaffold proteins: hubs for controlling the flow of cellular information. Science 332, 680–686, doi:10.1126/science.1198701 (2011).

2 Perez, A. M. et al. A Localized Complex of Two Protein Oligomers Controls the Orientation of Cell Polarity. mBio 8, doi:10.1128/mBio.02238-16 (2017).

3 Lasker, K., Mann, T. H. & Shapiro, L. An intracellular compass spatially coordinates cell cycle modules in Caulobacter crescentus. Current opinion in microbiology 33, 131–139, doi:10.1016/j.mib.2016.06.007 (2016).

4 Curtis, P. D. & Brun, Y. V. Getting in the loop: regulation of development in Caulobacter crescentus. Microbiology and molecular biology reviews : MMBR 74, 13–41, doi:10.1128/MMBR.00040-09 (2010).

5 Bergé, M. & Viollier, P. H. End-in-Sight: Cell Polarization by the Polygamic Organizer PopZ. Trends in microbiology 26, 363–375, doi: https://doi.org/10.1016/j.tim.2017.11.007 (2018).

6 Matroule, J. Y., Lam, H., Burnette, D. T. & Jacobs-Wagner, C. Cytokinesis monitoring during development; rapid pole-to-pole shuttling of a signaling protein by localized kinase and phosphatase in Caulobacter. Cell 118, 579–590, doi:10.1016/j.cell.2004.08.019 (2004).

7 Wheeler, R. T. & Shapiro, L. Differential localization of two histidine kinases controlling bacterial cell differentiation. Molecular cell 4, 683–694 (1999).

8 Jacobs, C., Hung, D. & Shapiro, L. Dynamic localization of a cytoplasmic signal transduction response regulator controls morphogenesis during the Caulobacter cell cycle. Proceedings of the National Academy of Sciences of the United States of America 98, 4095–4100, doi:10.1073/pnas.051609998 (2001).

9 Angelastro, P. S., Sliusarenko, O. & Jacobs-Wagner, C. Polar localization of the CckA histidine kinase and cell cycle periodicity of the essential master regulator CtrA in Caulobacter crescentus. Journal of bacteriology 192, 539–552, doi:10.1128/JB.00985-09 (2010).

10 Tsokos, C. G., Perchuk, B. S. & Laub, M. T. A dynamic complex of signaling proteins uses polar localization to regulate cell-fate asymmetry in Caulobacter crescentus. Developmental cell 20, 329–341, doi:10.1016/j.devcel.2011.01.007 (2011).

11 Tsokos, C. G., Perchuk, B. S. & Laub, M. T. A Dynamic Complex of Signaling Proteins Uses Polar Localization to Regulate Cell-Fate Asymmetry in Caulobacter crescentus. Developmental Cell 20, 329–341, doi:10.1016/j.devcel.2011.01.007 (2011).

12 Laub, M. T., Chen, S. L., Shapiro, L. & McAdams, H. H. Genes directly controlled by CtrA, a master regulator of the Caulobacter cell cycle. Proceedings of the National Academy of Sciences of the United States of America 99, 4632–4637, doi:10.1073/pnas.062065699 (2002).

13 Lasker, K. et al. Selective sequestration of signalling proteins in a membraneless organelle reinforces the spatial regulation of asymmetry in Caulobacter crescentus. Nature microbiology 5, 418–429, doi:10.1038/s41564-019-0647-7 (2020).

14 Laub, M. T., Chen, S. L., Shapiro, L. & McAdams, H. H. Genes directly controlled by CtrA, a master regulator of the Caulobacter cell cycle. Proceedings of the National Academy of Sciences of the United States of America 99, 4632–4637, doi:10.1073/pnas.062065699 (2002).

15 Bowman, G. R. et al. A polymeric protein anchors the chromosomal origin/ParB complex at a bacterial cell pole. Cell 134, 945–955, doi:10.1016/j.cell.2008.07.015 (2008).

16 Holmes, J. A. et al. Caulobacter PopZ forms an intrinsically disordered hub in organizing bacterial cell poles. Proceedings of the National Academy of Sciences of the United States of America 113, 12490–12495, doi:10.1073/pnas.1602380113 (2016).

17 Ptacin, J. L. et al. Bacterial scaffold directs pole-specific centromere segregation. Proceedings of the National Academy of Sciences of the United States of America 111, E2046–2055, doi:10.1073/pnas.1405188111 (2014).

18 Holmes, J. A. et al. Caulobacter PopZ forms an intrinsically disordered hub in organizing bacterial cell poles. Proceedings of the National Academy of Sciences, doi:10.1073/pnas.1602380113 (2016).

19 Viollier, P. H., Sternheim, N. & Shapiro, L. Identification of a localization factor for the polar positioning of bacterial structural and regulatory proteins. Proceedings of the National Academy of Sciences of the United States of America 99, 13831–13836, doi:10.1073/pnas.182411999 (2002).

20 Hinz, A. J., Larson, D. E., Smith, C. S. & Brun, Y. V. The Caulobacter crescentus polar organelle development protein PodJ is differentially localized and is required for polar targeting of the PleC development regulator. Molecular microbiology 47, 929–941, doi:10.1046/j.1365-2958.2003.03349.x (2003).

21 Curtis, P. D. et al. The scaffolding and signalling functions of a localization factor impact polar development. Molecular microbiology 84, 712–735, doi:10.1111/j.1365-2958.2012.08055.x (2012).

22 Lawler, M. L., Larson, D. E., Hinz, A. J., Klein, D. & Brun, Y. V. Dissection of functional domains of the polar localization factor PodJ in Caulobacter crescentus. Molecular microbiology 59, 301–316, doi:10.1111/j.1365-2958.2005.04935.x (2006).

23 Bowman, G. R. et al. Caulobacter PopZ forms a polar subdomain dictating sequential changes in pole composition and function. Molecular microbiology 76, 173–189, doi:10.1111/j.1365-2958.2010.07088.x (2010).

24 Schrader, J. M. et al. Dynamic translation regulation in Caulobacter cell cycle control. Proceedings of the National Academy of Sciences of the United States of America 113, E6859–E6867, doi:10.1073/pnas.1614795113 (2016).

25 Balleza, E., Kim, J. M. & Cluzel, P. Systematic characterization of maturation time of fluorescent proteins in living cells. Nature methods 15, 47–51, doi:10.1038/nmeth.4509 (2018).

26 Schrader, J. M. et al. Dynamic translation regulation in Caulobacter cell cycle control. Proceedings of the National Academy of Sciences 113, E6859–E6867, doi:10.1073/pnas.1614795113 (2016).

27 Ebersbach, G., Briegel, A., Jensen, G. J. & Jacobs-Wagner, C. A self-associating protein critical for chromosome attachment, division, and polar organization in caulobacter. Cell 134, 956–968, doi:10.1016/j.cell.2008.07.016 (2008).

28 Berge, M. et al. Modularity and determinants of a (bi-)polarization control system from free-living and obligate intracellular bacteria. Elife 5, doi:10.7554/eLife.20640 (2016).

29 Lam, H., Schofield, W. B. & Jacobs-Wagner, C. A landmark protein essential for establishing and perpetuating the polarity of a bacterial cell. Cell 124, 1011–1023, doi:10.1016/j.cell.2005.12.040 (2006).

30 Laloux, G. & Jacobs-Wagner, C. Spatiotemporal control of PopZ localization through cell cycle-coupled multimerization. The Journal of cell biology 201, 827–841, doi:10.1083/jcb.201303036 (2013).

31 Duerig, A. et al. Second messenger-mediated spatiotemporal control of protein degradation regulates bacterial cell cycle progression. Genes & development 23, 93–104, doi:10.1101/gad.502409 (2009).

32 Chen, J. C., Viollier, P. H. & Shapiro, L. A membrane metalloprotease participates in the sequential degradation of a Caulobacter polarity determinant. Molecular microbiology 55, 1085–1103, doi:MMI4443 [pii] 10.1111/j.1365-2958.2004.04443.x [doi] (2005).

33 Schofield, W. B., Lim, H. C. & Jacobs-Wagner, C. Cell cycle coordination and regulation of bacterial chromosome segregation dynamics by polarly localized proteins. The EMBO journal 29, 3068–3081, doi:10.1038/emboj.2010.207 (2010).

34 Ptacin, J. L. et al. A spindle-like apparatus guides bacterial chromosome segregation. Nature cell biology 12, 791–U746, doi:10.1038/ncb2083 (2010).

35 Thanbichler, M. & Shapiro, L. MipZ, a spatial regulator coordinating chromosome segregation with cell division in Caulobacter. Cell 126, 147–162, doi:10.1016/j.cell.2006.05.038 (2006).

36 Chen, J. C. et al. Cytokinesis signals truncation of the PodJ polarity factor by a cell cycle-regulated protease. The EMBO journal 25, 377–386, doi:10.1038/sj.emboj.7600935 (2006).

37 Anderson-Furgeson, J. C., Zupan, J. R., Grangeon, R. & Zambryski, P. C. Loss of PodJ in Agrobacterium tumefaciens Leads to Ectopic Polar Growth, Branching, and Reduced Cell Division. Journal of bacteriology 198, 1883–1891, doi:10.1128/jb.00198-16 (2016).

38 Grangeon, R., Zupan, J., Jeon, Y. & Zambryski, P. C. Loss of PopZ At activity in Agrobacterium tumefaciens by Deletion or Depletion Leads to Multiple Growth Poles, Minicells, and Growth Defects. mBio 8, doi:10.1128/mBio.01881-17 (2017).

39 Radhakrishnan, S. K., Thanbichler, M. & Viollier, P. H. The dynamic interplay between a cell fate determinant and a lysozyme homolog drives the asymmetric division cycle of Caulobacter crescentus. Genes & development 22, 212–225, doi:10.1101/gad.1601808 (2008).

40 Viollier, P. H., Sternheim, N. & Shapiro, L. A dynamically localized histidine kinase controls the asymmetric distribution of polar pili proteins. The EMBO journal 21, 4420–4428, doi:10.1093/emboj/cdf454 (2002).

41 Al-Husini, N., Tomares, D. T., Childers, W. S. & Schrader, J. α-proteobacterial RNA degradosomes assemble liquid-liquid phase separated RNP bodies. Molecular cell (2018).

42 Al-Husini, N. et al. BR-Bodies Provide Selectively Permeable Condensates that Stimulate mRNA Decay and Prevent Release of Decay Intermediates. Molecular cell, doi:10.1016/j.molcel.2020.04.001 (2020).

43 Monterroso, B. et al. Bacterial FtsZ protein forms phase-separated condensates with its nucleoid-associated inhibitor SlmA. EMBO reports 20, doi:10.15252/embr.201845946 (2019).

44 Heinkel, F. et al. Phase separation and clustering of an ABC transporter in Mycobacterium tuberculosis. Proceedings of the National Academy of Sciences of the United States of America 116, 16326–16331, doi:10.1073/pnas.1820683116 (2019).

45 Lasker, K. et al. Selective sequestration of signalling proteins in a membraneless organelle reinforces the spatial regulation of asymmetry in Caulobacter crescentus. Nature microbiology, doi:10.1038/s41564-019-0647-7 (2020).

46 Tropini, C. & Huang, K. C. Interplay between the localization and kinetics of phosphorylation in flagellar pole development of the bacterium Caulobacter crescentus. PLoS computational biology 8, e1002602, doi:10.1371/journal.pcbi.1002602 (2012).

47 Ely, B. Genetics of Caulobacter crescentus. Methods in enzymology 204, 372–384 (1991).

48 Evinger, M. & Agabian, N. Envelope-associated nucleoid from Caulobacter crescentus stalked and swarmer cells. Journal of bacteriology 132, 294–301 (1977).

49 Ducret, A., Quardokus, E. M. & Brun, Y. V. MicrobeJ, a tool for high throughput bacterial cell detection and quantitative analysis. Nature microbiology 1, 16077, doi:10.1038/nmicrobiol.2016.77 (2016).

50 Schindelin, J. et al. Fiji: an open-source platform for biological-image analysis. Nature methods 9, 676–682, doi:10.1038/nmeth.2019 (2012).

51 Preibisch, S., Saalfeld, S. & Tomancak, P. Globally optimal stitching of tiled 3D microscopic image acquisitions. Bioinformatics 25, 1463–1465, doi:10.1093/bioinformatics/btp184 (2009).

